# The Northern Colours: Isolation and Characterisation of 4 Pigment-Producing Bacteria from the Arctic

**DOI:** 10.1101/2024.02.13.579883

**Authors:** Jenifar Das, Ashish Kumar Singh

## Abstract

Due to the adverse effects of synthetic colours on human health and the environment, there is a rapid shift towards the use of colours from natural sources like plants and microorganisms. Many pigment-producing microorganisms are identified and isolated from extreme environments like glaciers, ice cores, marine surface waters, etc. In this study, we have isolated 4 distinct pigment-producing bacterial strains from an Arctic stone sample collected from the vicinity of the Indian Research Station *Himadri* (78°55^′^*N* 11°56^′^*E*), located at the International Arctic Research Base, NyÅlesund, Svalbard, Norway. Pigment production was optimised by identifying the right growth medium, temperature, pH, and incubation period. The morphological, cultural, and biochemical characteristics were identified using several experiments like Gram Staining, Catalase Test, Oxydative-Fermentative Test, etc. The objective of this study is to identify novel bacterial strains capable of producing distinct pigments for pharmaceutical and industrial applications.

## 1. Introduction

The use of artificial colours is commonplace in the food, pharmaceutical, cosmetic, and textile industries. Their production is easy and low-cost yet comes with the advantages of high colouring strength, attractive hues, and good stability (Scotter, 2011). However, as most of the artificial colours are produced by chemical synthesis, they have adverse effects on both, human health and the environment (Sen et al., 2019). For example, several Food and Drug Administration (FDA) approved artificial colours were later found to be carcinogenic (Narsing Rao et al., 2017). Thanks to the ever-growing awareness related to health and the environment, there is a rapid shift towards the use of colours from natural sources like plants and microorganisms (Cavalcante et al., 2023; El-Sayed et al., 2022; Aman Mohammadi et al., 2022; Lagashetti et al., 2019). Natural colours are harmless, biodegradable, and noncarcinogenic (Cristea and Vilarem, 2006). The market intelligence firm *Grand View Research, Inc*. estimates that the natural food colour market size will reach $2.52 billion by the year 2030.

Natural colours are basically pigments produced by different plants and microorganisms for their survival. For example, plants use pigments for photosynthesis (Siefermann-Harms, 1990), pollination (Choo, 2018), protection against UV and visible light (Tanaka et al., 2008), etc. Similarly, microorganisms use pigments as a source of energy (Madigan et al., 2012), as an anti-oxidant (Wada et al., 2013), to mitigate stress (Martín-Cerezo et al., 2015), etc. Hence, in addition to colour, natural pigments also offer antimicrobial, antioxidant, anti-inflammatory, anti-obesity, and anti-cancer properties (Tuli et al., 2015). Microorganism-based natural pigments are preferred over their plant-based counterparts as they have better solubility, stability, and scalability (Narsing Rao et al., 2017). Among microorganisms, algae, bacteria, and fungi are the major sources of natural pigments (Ramesh et al., 2019; Pandey et al., 2018; Narsing Rao et al., 2017; Choi et al., 2015; Joshi et al., 2003). They can produce natural pigments like carotenoids, melanins, flavins, quinines, monascins, violancein, etc (Dufossé, 2006). However, to meet the increasing market demand for natural colours, it is imperative to identify more and more pigment-producing microorganisms (Sajjad et al., 2020).

Many pigment-producing microorganisms are identified and isolated from extreme environments like glaciers, ice cores, marine surface waters, etc (Agogué et al., 2005; Foght et al., 2004). The Arctic zone has such an extreme environment and is home to many psychrophilic microorganisms that are yet to be explored. To survive in that environment with low temperatures, solar radiation, higher salinity, desiccation, and repeated freeze-thaw cycles, psychrophiles undergo morphology and genome modification (Uddin et al., 2022). They produce secondary metabolites, such as pigments, to reduce solar radiation damage. Pigments also help in osmoregulation to tolerate extreme salinity, desiccation, and freeze-thaw cycles (Mueller et al., 2005). If necessary, pigments can act as antimicrobials for protection against neighbouring psychrophiles or as shields to evade antibiosis. Thus, psychrophiles have become one of the most promising sources of natural pigments. Multiple field works in the Arctic zone have reported high bacterial abundance in soil, sediments, permafrost, ice, and water (Singh et al., 2014; Forschner et al., 2009; Hansen et al., 2007).

In this study, we have examined an Arctic stone sample with pigmented spots to identify potentially pigment-producing bacteria. We were able to isolate 4 distinct pigment-producing bacterial strains. We optimised the pigment production by identifying the right growth medium, temperature, pH, and incubation period. The morphological, cultural, and biochemical characteristics were then identified using several experiments.

## 2. Materials and Methods

### 2.1. Sampling Site

The sample was collected from the vicinity of the Indian Research Station *Himadri* (78°55^′^*N* 11°56^′^*E*), located at the International Arctic Research Base, NyÅlesund, Svalbard, Norway. It is located at a distance of 1,200 kilometres from the North Pole. The sample was collected in a sterile plastic bag and transported to the laboratory in an icebox at ≤4°C.

### 2.2. Isolation of Pigment-Producing Bacteria

The Arctic stone sample, as shown in Figure 1, is finely crushed to identify all the bacteria in it. The crushed sample is then diluted up to 10^−7^ using autoclaved distilled water. From the 10^−7^ dilutions, 0.1 ml each was spread into solid plates of *Luria Agar (LA)* (Himedia, India, pH 7.0) and *Nutrient Agar (NA)* (Himedia, India, pH 7.4). The inoculated plates were incubated at 25°C and 37°C for 24-72 hours until single bacterial colonies appeared. As we were interested in identifying pigment-producing bacteria, after streaking to purity on NA, we could isolate 4 distinct pigment-producing bacterial strains.

**Figure 1:**
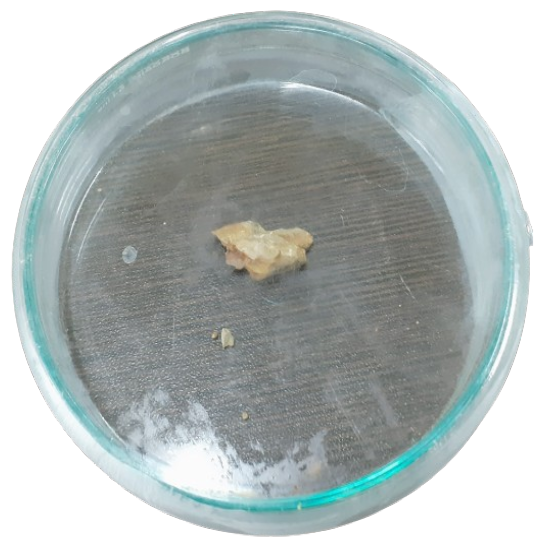
Arctic stone sample.

### 2.3. Characterisation of Pigment-Producing Bacteria

#### 2.3.1. Cultural Characteristics

Gram-staining properties were investigated after incubating each culture in NA for 24-72 hours. One drop of a suspended culture was transferred on a slide with an inoculation loop to prepare a slide smear. The culture was evenly spread over a circle of 1.5 cm in diameter. The slide was allowed to air dry and fixed over a gentle flame. The slide was stained with *Crystal Violet* for 60 seconds and rinsed with distilled water. The slide was stained again with Iodine solution for 30 seconds and rinsed. Then it was decolourised, counterstained again with *Safranin* for 30 seconds, and rinsed. The dried slide is examined under a microscope. It was repeated for other cultures.

#### 2.3.2. Biochemical Characteristics

Each isolated strain is tested for its *Catalase, Oxidase, Urease, Indole* production abilities, and *Citrate* utilisation abilities. The *Oxidative-Fermentative (OF)* test was also conducted for each strain using Hugh and Leifson’s OF basal medium.

### 2.4. Optimisation of Time and pH for Pigment Production

Pigment production was optimised for each culture in NA with varying incubation periods (24-72 hours) and pHs (5-9).

## 3. Results and Discussion

### 3.1. Isolation of Pigment-Producing Bacterial Strains

We have found multiple strains of microorganisms on the examined Arctic stone sample. Between the LA and NA mediums we used to culture the strains, the most diversities in colony morphology were found in NA. Among the different dilutions we have used to culture the strains, the most vivid colonies started to appear from 10^−4^, as shown in Figure 2. As this study focused on pigment-producing bacteria, we isolated 4 strains with distinct pigments, as shown in Figure 3. *Strain 1, Strain 2, Strain 3* and *Strain 4* produces *Yellow, Orange-Red, Pink* and *Creamy* pigments respectively. While each strain has a different incubation period between 24-72 hours, all of them have the optimum growth at 25°C. pH is another important factor for stimulating growth, and we have observed that none of the strains had any growth till pH 5.0. All of them are found to be alkaline in nature, showing full growth only from pH 6.0 onwards.

**Figure 2:**
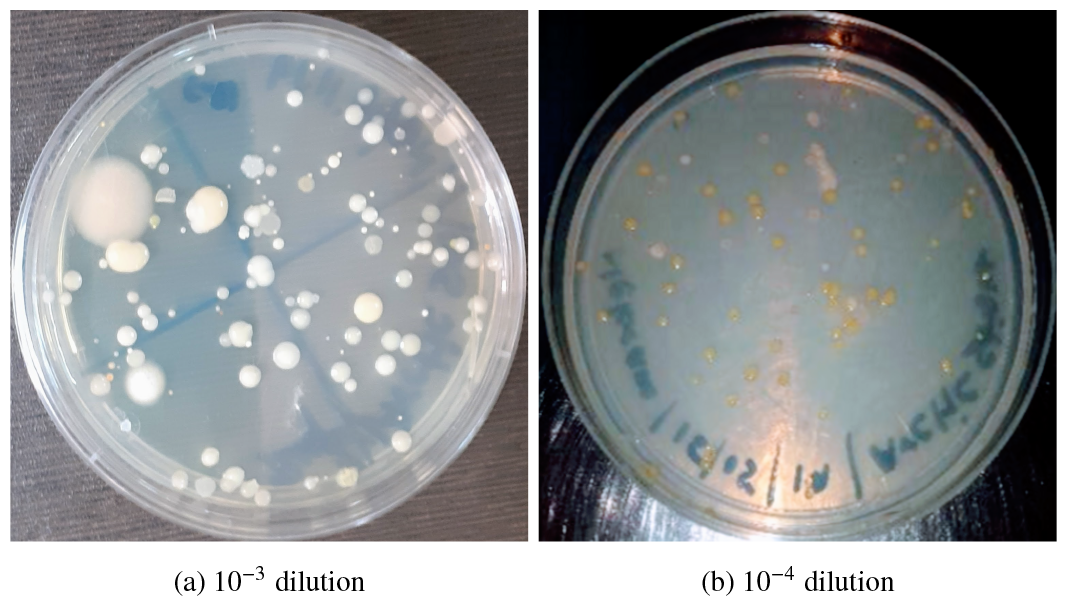
Bacterial culture plate in NA medium

**Figure 3:**
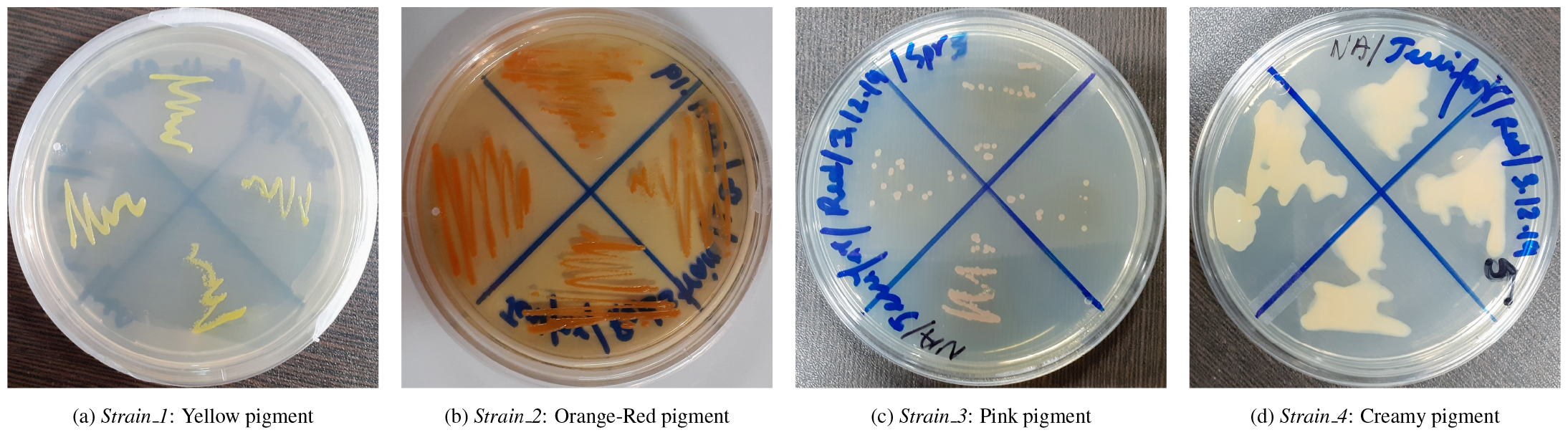
Isolated bacterial strains

### 3.2. Characterisation of Pigment-Producing Bacterial Strains

Table 1 presents all the characteristics of the isolated bacterial strains, where the columns are colour-coordinated with the pigment produced by the respective strains. Morphological observation under the microscope showed that all the strains had spherical shapes, with *Strain 1* and *Strain 3* having coccus shape, while *Strain 2* and *Strain 4* having clusters of coccus (streptococci) shape. Gram staining experiment revealed that only *Strain 2* is gram-positive, and all others are gram-negative. A gram-positive strain will have a purple stain, as shown in Figure 4b, indicating a thick peptidoglycan layer in the cell wall.

**Table 1:**
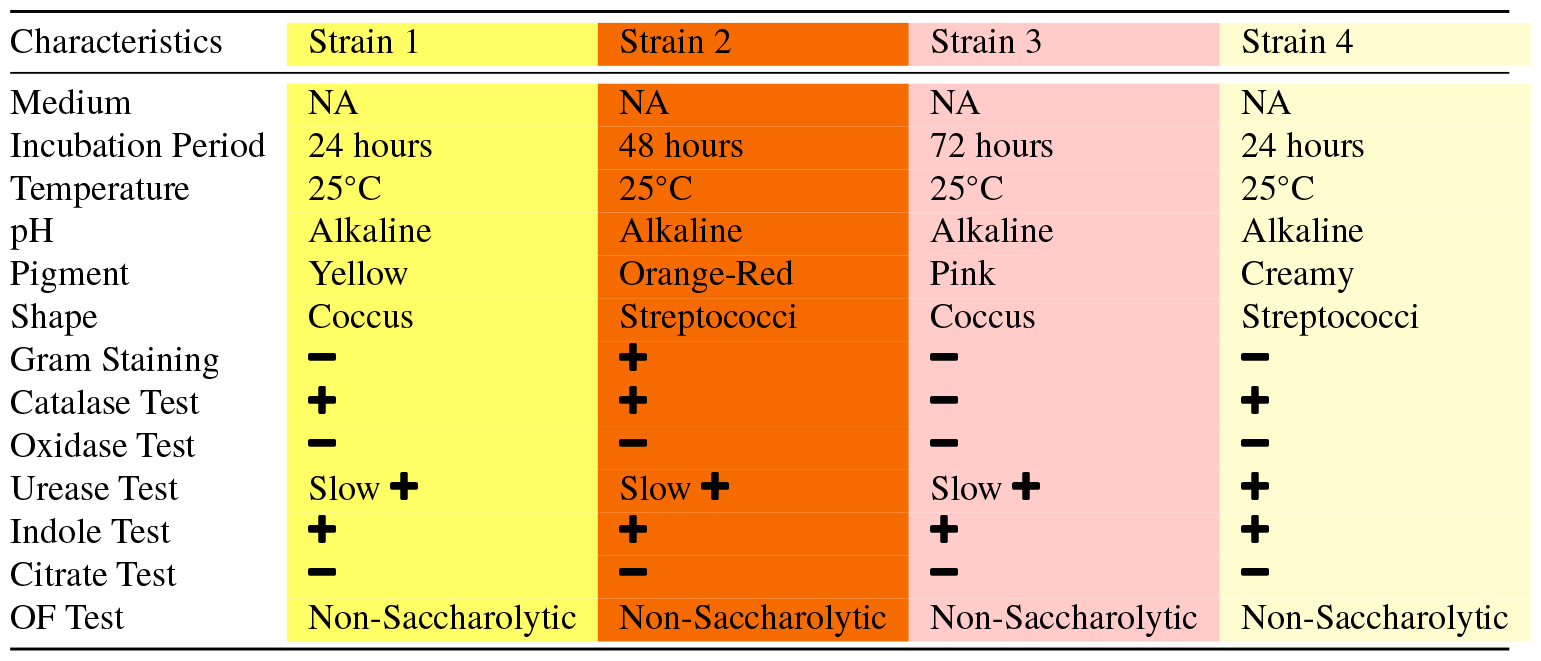
Characteristics of the isolated pigment-producing bacterial strains.

**Figure 4:**
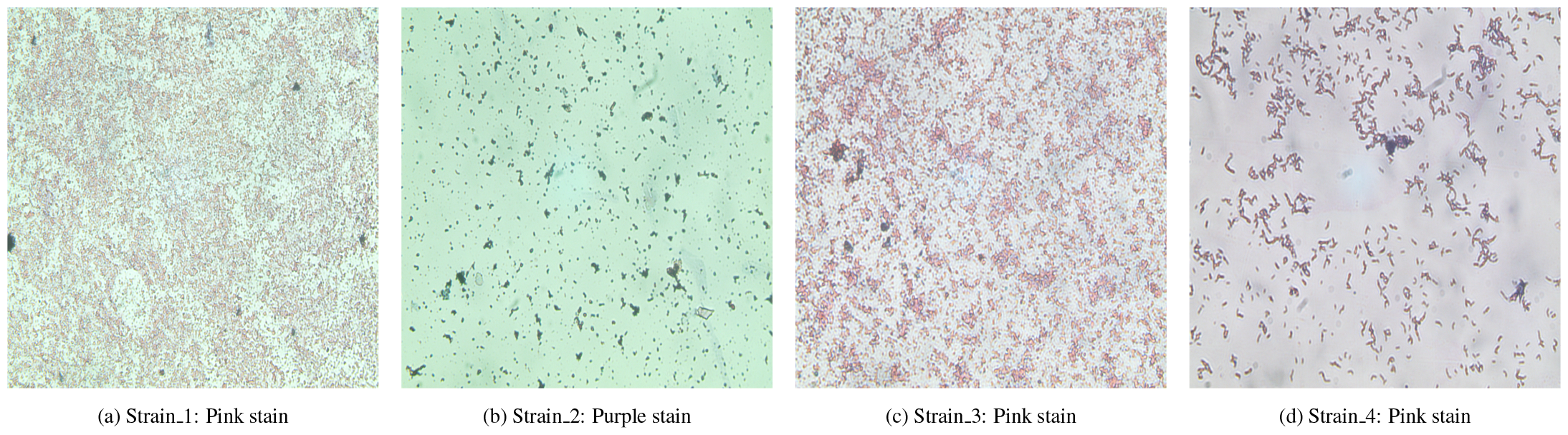
The Gram Staining experiment

Whereas a gram-negative strain will have a pink or red stain, indicating a thinner peptidoglycan layer in the cell wall. Thus, according to Bergey’s Manual (Bergey, 1994), *Strain 1, Strain 3* and *Strain 4* could belong to *Veillonella* or *Neisseria* genus. Whereas *Strain 1* could belong to *Stephylococcus* genus.

*Catalase* is a common enzyme found in bacteria and other organisms exposed to oxygen. It helps organisms protect their cells from oxidative damage by catalysing the decomposition of hydrogen peroxide to water and oxygen. Our test found that *Strain 1, Strain 2* and *Strain 4* are catalase-positive while *Strain 3* is catalase-negative. *Oxidase* is another enzyme that catalyses the oxidation-reduction (redox) reaction. Our test found that all the strains are oxidase-negative. *Urease* is another enzyme that catalyses the hydrolysis of urea into ammonia and carbon dioxide. Our test found that all the strains are urease-positive with *Strain 1, Strain 2* and *Strain 3* being slow.

The *Indole Test* is performed to determine the ability of a strain to convert tryptophan into indole. Our test found that all the isolated bacterial strains were indole-positive. The *Citrate Test* is performed to determine the ability of a strain to utilise citrate as its carbon and energy source. Our test found that all the isolated bacterial strains were citrate-negative. The *Oxydative-Fermentative (OF) Test* is performed to determine if and how a strain metabolites carbohydrate (glucose). Our test found that all the isolated bacterial strains were *Non-Saccharolytic*, meaning none of them could metabolise glucose.

### 3.3. Special Observation from Strain 3

For all the experiments till this point, *Strain 3* had Pink pigment after 72 hours of incubation at 25°C. However, during characterisation we observed that after an additional 24-48 hours of the incubation period, *Strain 3* starts producing *Brown* pigment. As shown in Figure 5b, brown pigments are produced at the edges of the culture plate after 72+48 hours of incubation. As shown in Figure 6b, a similar modification from pink to brown pigment production was observed on the culture flask. Once brown, there was no changing back to the pink pigment.

**Figure 5:**
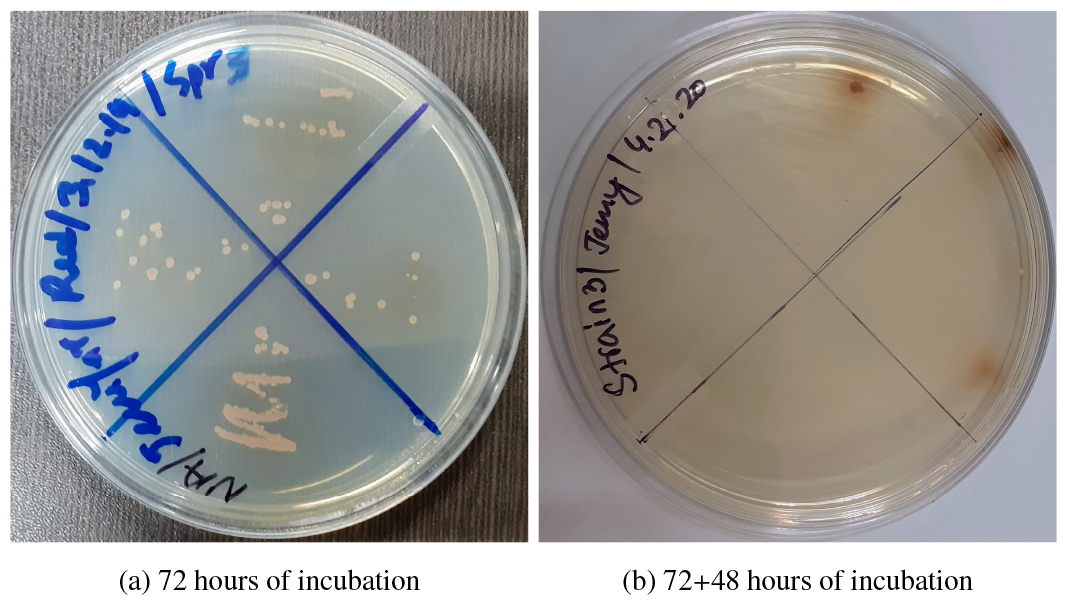
Culture plate of Strain 3 after different incubation periods

**Figure 6:**
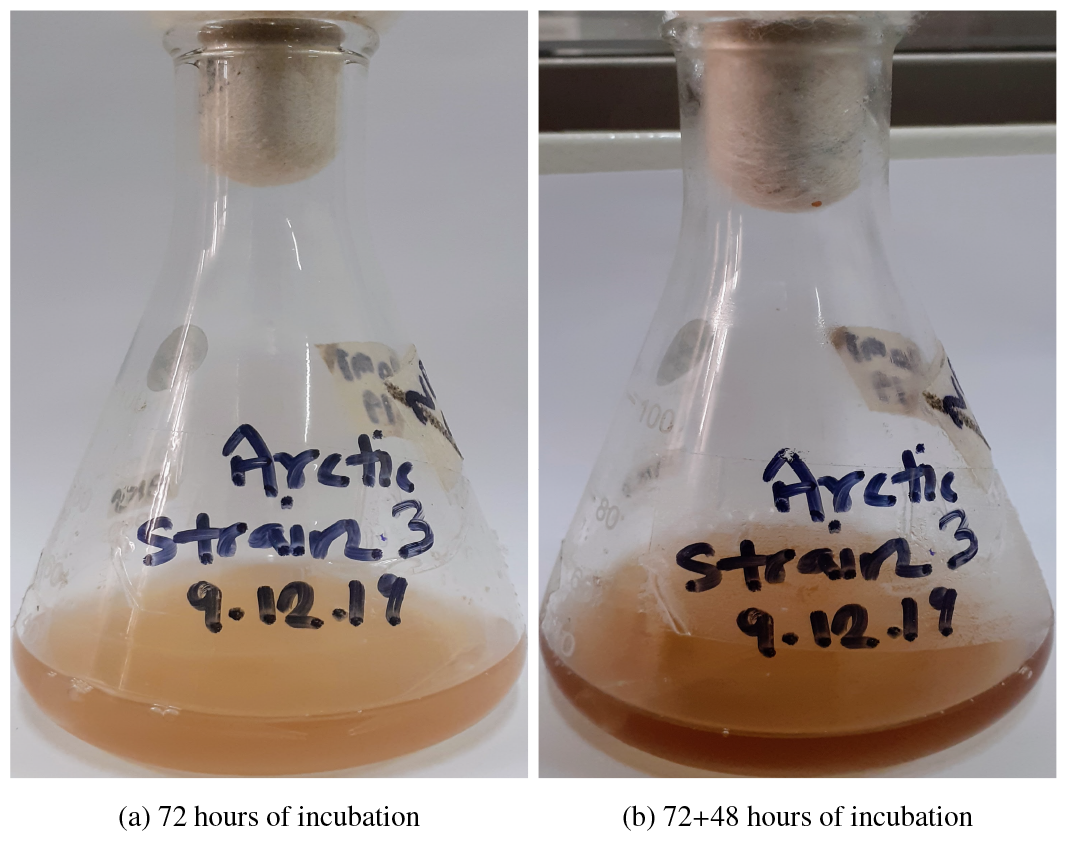
Culture flask of Strain 3 after different incubation periods

## 4. Conclusion

In this study, we have isolated 4 distinct pigment-producing bacterial strains from an Arctic stone sample. Characterisation of the isolated strains shows great potential for medical and industrial applications. Optimal pigment production is achieved at 25°C, which means that these strains can be easily replicated at a low temperature. Hence, these isolated strains will have lower production and maintenance costs thereby making a strong case for their adoption. The capability of *Strain 3* producing two different pigments can exploited as necessary.

